# Cue-reactivity in the ventral striatum characterizes heavy cannabis use, whereas reactivity in the dorsal striatum mediates dependent use

**DOI:** 10.1101/516385

**Authors:** Xinqi Zhou, Kaeli Zimmermann, Fei Xin, Weihua Zhao, Roelinka Derckx, Anja Sassmannshausen, Dirk Scheele, Rene Hurlemann, Bernd Weber, Keith M. Kendrick, Benjamin Becker

## Abstract

**Background:** Animal models of addiction suggest that the transition from incentive-driven to habitual and ultimately compulsive drug use is mediated by a shift from ventral to dorsal striatal cue-control over drug seeking. Previous studies in human cannabis users reported elevated trait impulsivity and cue-reactivity in striatal circuits, however, these studies were not able to separate addiction-related from exposure-related adaptations.

**Methods:** To differentiate the adaptive changes, the present functional magnetic resonance imaging study examined behavioral and neural cue-reactivity in dependent (n = 18) and non-dependent (n = 20) heavy cannabis users and a non-using reference group (n = 44).

**Results:** Irrespective of dependence status, cannabis users demonstrated elevated trait impulsivity as well as increased ventral striatal reactivity and striato-frontal coupling in response to drug cues. Dependent users selectively exhibited dorsal-striatal reactivity and decreased striato-limbic coupling during cue-exposure. An exploratory analysis revealed that higher ventral caudate cue-reactivity was associated with stronger cue-induced arousal and craving in dependent users, whereas this pattern was reversed in non-dependent users.

**Conclusions:** Together the present findings suggest that an incentive sensitization of the ventral striatal reward system may promote excessive drug use in humans, whereas adaptations in dorsal striatal systems engaged in habit formation may promote the transition to addictive use.

## Introduction

Drug addiction is a chronically relapsing disorder of the brain characterized by compulsive drug use, loss of behavioral control and an intense, overwhelming desire to consume the drug during deprivation or exposure to drug-associated cues (craving) (1). The transition from volitional to compulsive use is accompanied by dysregulations in the brain’s motivational circuitry. Neuroplastic changes promote exaggerated incentive salience and compulsive-like habitual responses to the drug itself and cues repeatedly paired with the drug. Compelling evidence from animal models suggests that these cues acquire excessive motivational significance via drug-induced dysregulations in operant and instrumental learning mechanisms (2, 3). The striatal dopamine system critically contributes to the acute reinforcing effects across different classes of drugs (4) as well as operant and instrumental learning mechanism engaged in incentive salience and habit formation (3, 5, 6). The multi-facetted contribution of the striatum to the development and maintenance of addiction is mirrored in the segregated circuits and heterogenous functional organization of this structure. Ventral striatal regions, connected to limbic and orbitofrontal regions, are engaged in reward processing, incentive-based learning and impulsive behavior, whereas dorsal regions contribute to habit formation, compulsive behavior and regulatory control via connections with ventral striatal and dorsomedial prefrontal regions (7–10). In line with this functional differentiation, animal models suggest that the transition from incentive-driven to compulsive drug use is mediated by a shift from ventral-to dorsal-striatal control over behavior (2).

Cannabis is the most widely used illicit drug worldwide, with more than 2% of the world’s population consuming cannabis on a regular basis (UNODC, https://www.unodc.org/wdr2018/). Similar to other potentially addictive substances, cannabis leads to an acute increase in striatal dopamine transmission (11)(however see also (12)). Accumulating evidence suggests that frequent cannabis use is accompanied by brain structural and functional maladaptations in fronto-striato-limbic circuits engaged in reward, learning and cognitive control which may mediate the transition to addiction (13–17). However, even with heavy use less than 40% of cannabis users will develop addictive patterns of use over the course of three years (18). Given that the majority of previous studies did not consider the dependence status of cannabis users, neuroplastic alterations directly related to addictive behavior cannot be disentangled from adaptations related to cannabis exposure or compensatory neuroadaptations (14). Against this background, a recent review proposed that comparing dependent and non-dependent cannabis users represents a crucial next step to delineate neuroadaptations specifically related to cannabis addiction while controlling for exposure related adaptations (15).

This approach has been successfully employed to demonstrate brain structural changes in orbitofrontal and hippocampal regions. However, as yet, it is still elusive whether activations in striatal subregions differentiate dependent from non-dependent users and thus may specifically characterize addictive patterns of use (19–21). An initial fMRI cue-reactivity study reported increased functional crosstalk between the ventral striatum and reward and salience-processing core hubs such as the amygdala in dependent relative to non-dependent users (22). However, these findings remain inconclusive due to the lack of a non-using reference group, as well as differences in cannabis craving and nicotine use between the cannabis using groups which may affect neural cue-reactivity (see also cross-cue reactivity in (23, 24)).

Importantly, drug cue-reactivity is thought to reflect incentive motivation as well as compulsive processes in addiction (25) and is mediated by exaggerated striatal reactivity towards drug-associated stimuli across addictive disorders (26–29). Converging evidence from previous studies demonstrated cue-induced exaggerated reactivity in dopaminergic reward pathways such as the ventral tegmental area (VTA) in frequent cannabis users (30, 31). However, exaggerated VTA reactivity may reflect neuroplastic processes as a consequence of chronic cannabis exposure (30), whereas associations between higher cannabis use problems and striatal reactivity in the context of a lack of associations with exposure-related indices suggests that neuroadaptations in the striatum may mediate the addictive process (30, 31). Further support for the clinical relevance of dorsal striatal alterations comes from a prospective study reporting that cue-reactivity in this region predicts severity of cannabis use problems over three years (32) (convergent findings in alcohol addiction see (33)). Likewise, translational studies found a distinct involvement of the dorsal striatum in compulsive drug use in rodents (34) and humans (35). However, despite increasing treatment demands for cannabis use disorder (UNODC, https://www.unodc.org/wdr2018/) and comprehensive animal models suggesting a ventral to dorsal striatal shift associated with drug cue-controlled behavior, specific contributions of striatal neuroadaptations to cannabis addiction in humans remains to be determined.

Against this background, the present fMRI study examined neural cue-reactivity in dependent and non-dependent cannabis users as well as a carefully matched reference group. Based on comprehensive animal models of addiction (2, 3), we expected that both groups of cannabis users would exhibit exaggerated cue reactivity in the ventral striatum, whereas only dependent users would exhibit exaggerated reactivity in the dorsal striatum. Furthermore, impulsivity represents a trait-endophenotype marker for escalating substance use (36, 37)(but see (38, 39)) and is specifically mediated by ventral – but not dorsal – striatal regions (40, 41). Therefore we hypothesized that both groups of cannabis users would report increased levels of trait-impulsivity.

## Materials and Methods

### Participants and procedures

A total of n = 51 male cannabis users and n = 52 matched non-using controls were recruited. In order to disentangle dependence and exposure-related adaptations, the cannabis group was stratified according to dependence status determined by a structured clinical interview according to DSM-IV criteria (n = 26 cannabis users fulfilled; n = 25 cannabis users did not fulfill criteria for cannabis dependence; Mini-International Neuropsychiatric Interview DSM-IV, MINI (42)). To reduce variance due to sex- and menstrual cycle-dependent effects on striatal activity (43) and craving (44) only males were enrolled. Further study criteria aimed at reducing potential confounding effects of other factors such as co-morbid disorders or life-time and current co-use of other substances (details see **Supplemental Material**). Eligible participants underwent an assessment of additional confounders, trait impulsivity (as assessed by the Barrat Impulsivness Scale, BIS) and a validated fMRI blocked-design cue-reactivity paradigm during which cannabis and neutral stimuli were presented. To assess behavioral indices of cue-reactivity participants were required to rate their cannabis craving (before and after fMRI) and arousal-induction by the stimuli (during fMRI). Following initial quality assessments of the data, n = 9 cannabis users (n = 4 dependent, n = 5 non-dependent) and n = 8 controls were excluded due to low MRI data quality or excessive head motion (> 3 mm or > 3°). In addition, n = 4 dependent cannabis users were excluded due to exceeding the cut-off for co-use of other drugs. Consequently, n = 18 dependent cannabis users, n = 20 non-dependent cannabis users and n = 44 healthy controls were included in the final analysis (detailed procedures are given in **Supplemental Material**, for sample details see **Table 1).**

**Table 1.** Group characteristics and drug use parameters

| Measure | Dependent<br>M (SD) | Non-dependent<br>M (SD) | Controls<br>M (SD) | F<br>($\chi^2$ / t) | p |
| --- | --- | --- | --- | --- | --- |
| Age (years) | 22.94 (2.71) | 21.48 (2.54) | 23.2 (4.32) | 1.59 | 0.211 |
| Education (years) | 15.33 (2.46) | 14.35 (1.73) | 15.61 (2.46) | 2.08 | 0.132 |
| Attention (D2) | 183 (39.19) | 179 (35.54) | 197 (39.79) | 1.86 | 0.163 |
| State anxiety (STAI) | 34.83 (7.86) | 32.85 (4.93) | 31.91 (4.75) | 1.74 | 0.182 |
| Number of smokers | 17 | 14 | 33 | 3.82 | 0.148<sup>a</sup> |
| Age of first nicotine use | 15.59 (2.62) | 16.54 (2.1) | 16.06 (1.82) | 0.77 | 0.466 |
| Years of nicotine use | 6.21 (3.12) | 5.07 (2.95) | 5.98 (3.81) | 0.47 | 0.629 |
| Cigarettes per day | 9.85 (6.16) | 9.24 (6.74) | 8.41 (5.11) | 0.37 | 0.693 |
| Pack-year | 3.32 (3.14) | 2.34 (1.83) | 2.92 (3.13) | 0.435 | 0.649 |
| Fagerstrom score | 2.24 (2.11) | 2.21 (1.76) | 1.58 (1.64) | 1.04 | 0.358 |
| Age of first alcohol intake | 15.75 (2.1) | 15.28 (1.8) | 15.83 (1.6) | 0.74 | 0.479 |
| Alcohol occasions per week<br>(day) | 1.36 (1.08) | 1.73 (1.16) | 1.05 (0.93) | 3.04 | 0.054 |
| Alcohol units per week | 10.13 (8.52) | 9.48 (7.28) | 8.6 (11.27) | 0.16 | 0.851 |
| Number - past ecstasy use | 11 | 10 | 4 |  |  |
| Lifetime occasions ecstasy | 8.36 (8.02) | 6.2 (6.09) | 2.25 (1.89) | 1.24 | 0.309 |
| Number - past cocaine use | 7 | 7 | 2 | - | - |
| Lifetime occasions cocaine | 2.57 (1.62) | 2.14 (1.07) | 5.5 (6.36) | 1.85 | 0.197 |
| Number past amphetam. use | 10 | 9 | 4 | - | - |
| Lifetime occasions amphetam. | 6 (4.74) | 3.5 (4.5) | 5.5 (5.2) | 0.7 | 0.508 |
| Number past hallucinog. use | 11 | 10 | - | - | - |
| Lifetime occasions hallucinog. | 5.36 (8.64) | 1.5 (0.97) | - | 1.47 | 0.171<sup>b</sup> |
| Number - past opiate use | 1 | 1 | - | - | - |
| Lifetime occasions opiate | 1 | 1 | - | - | - |
| Number past solvents use | 6 | 5 | - | - | - |
| Lifetime occasions solvents | 2 (0.89) | 3 (2) | - | - | - |
| Number past cannabis use | 18 | 20 | 33 | - | - |
| % Cannabis dependence | 100 | 0 | 0 | - | - |

Note: Number – past use refers to number of participants with lifetime experience of the respective substance; lifetime occasions refers to the mean lifetime occasions of the participants with experience of the respective drug, amphetam. abbreviates amphetamine, hallucinog. abbreviates hallucinogens.
<sup>a</sup> $\chi^2$ test, <sup>b</sup> independent sample t test

Participants were recruited in cooperation with drug counseling services in Germany. Written informed consent was obtained, procedures were approved by the local ethics committee (University of Bonn, Germany) and adhered to the latest revision of the Declaration of Helsinki.

### Analysis of between group differences in neural cue-reactivity

FMRI acquisition and processing details are provided in **Supplemental Material**. Group differences in neural cue reactivity (cannabis cue > neutral) were initially examined on the whole-brain level by means of a one-way ANOVA post-hoc t-tests comparing each cannabis group separately to the non-using controls (similar approach see (30, 45, 46)). An additional region-of-interest (ROI) analysis was conducted to specifically evaluate our a priori regional hypothesis on a differential involvement of the ventral versus dorsal striatum in cannabis addiction (see also (17)). To this end individual parameter estimates were extracted from atlas-based (47) ventral and dorsal striatal subregions. The ventral striatum (VS) encompassed ventral caudate and nucleus accumbens, the dorsal striatum (DS) combined the dorsal caudate, dorsolateral putamen and ventromedial putamen (36, 48, **Figure S2**). Extracted estimates were subjected to a mixed ANOVA model with the between-subject factor group (dependent versus non-dependent) and the within-subject factor subregion (ventral versus dorsal).

### Functional connectivity analysis

To further explore cannabis addiction-related alterations of ventral and dorsal striatal communication with other regions of the brain, task-modulated functional connectivity was computed (generalized context-dependent psychophysiological interactions, gPPI, analysis (49), CONN toolbox, http://www.nitrc.org/projects/conn, RRID: SCR_009550). Briefly, the VS and DS masks were employed as seed regions, experimental conditions were modelled in line with the BOLD level analysis and the cue-reactivity contrast (cannabis-cue > neutral) served as primary outcome. Network level alterations in cannabis users were determined by comparing the subregion-specific cue-reactivity connectivity maps between the groups by means of independent t test.

### Thresholding

Line with current recommendations for the application of cluster-based control of false-positives (50) the initial cluster-forming threshold was set to *p* < 0.001 (voxel-level, whole brain), and statistical significance was determined via cluster-level inference and family-wise error (FWE) control for multiple comparisons with *p*_*FWE*_ < 0.05. For post-hoc analyses appropriate Bonferroni-corrections were employed.

### Associations between behavioral and neural cue-reactivity

Given the importance of craving and drug-induced arousal in addiction (26, 51), associations between these variables and neural markers were explored (Pearson correlation). Given that most voxels differentiating the groups were located in the ventral caudate (both dependent group > control and non-dependent group > control, **Result**s, **Figure. 5A**) this region (vCa, from the brainetome atlas) was used to extract parameter estimates as an individual index of neural cue-reactivity. Group-specific associations between behavioral and neural cue reactivity were examined between vCa cannabis-cues responses and cue-induced arousal and craving.

## Results

### Potential confounding factors and cannabis use patterns

Age, education, attention, anxiety, nicotine and alcohol use were comparable between groups (**Table 1**). Both cannabis groups reported long-term, heavy cannabis use with comparable duration and lifetime amount. The groups reported comparable experience with other illicit drugs. However, dependent users reported earlier age at first use (mean age = 14.89, SD = 2.08) compared to non-dependent users (mean = 16.28, SD = 1.81) and a shorter time since last use (dependent, 39.78±32.95h; range: 24-168; non-dependent, 83.25±46.34h, range: 38-240). Consequently, these parameters were included as covariates in direct comparisons between the cannabis groups (details see **Table 2**).

**Table 2.** Cannabis use parameters

| Measure | Dependent<br>M (SD) | Non-dependent<br>M (SD) | T value | p |
| --- | --- | --- | --- | --- |
| Age of first cannabis use | 14.89 (2.08) | 16.28 (1.81) | 2.2 | 0.035 |
| Hours since last cannabis use | 39.78 (32.95) | 83.25 (46.34) | 3.3 | 0.002 |
| Duration of regular cannabis use (months) | 61.83 (34.61) | 55.38 (33.76) | 0.58 | 0.565 |
| Lifetime amount of cannabis (g) | 1583.27<br>(1191.93) | 984.45<br>(1023.68) | 1.67 | 0.104 |

### Between-group differences in trait impulsivity, cue-induced arousal and craving

Examining self-reported impulsivity (BIS total) revealed a significant main effect of group (*F*_(2,79)_=6.05, *p* = 0.004, *η*^*2*^ = 0.133), with post-hoc tests indicating that both cannabis groups reported significantly higher trait impulsivity than controls (non-dependent users > controls, *t* = 3.224, *p*_*bonferroni*_ = 0.006, Cohen’s *d* = 0.857, dependent users > controls, *t* = 2.214, *p*_*bonferroni*_ = 0.089, Cohen’s *d* = 0.653, see **Figure. 1A**) and no difference between cannabis groups (*p* > 0.44). Mixed ANOVA including the between-subject factor group (controls vs. non-dependent vs. dependent), the within-subject factor cue (neutral vs. cannabis) and arousal as dependent variable revealed significant main effects of both cue (*F*_(2,79)_ = 20.5, *p* < 0.001, *η*^*2*^ = 0.34) and group (*F*_(1,79)_ = 66.3, *p* < 0.001, *η*^*2*^ = 0.326) as well as a significant interaction (*F*_(2,79)_ = 29.1, *p* < 0.001, *η*^*2*^ = 0.286). Post-hoc tests revealed that arousal ratings for cannabis cues in both cannabis groups were significantly higher compared to controls (both *p*_*bonferroni*_ < 0.001). In addition, in both cannabis groups cannabis cues were rated as more arousing than neutral stimuli (both *p*_*bonferroni*_ < 0.001). Importantly, there were no between-group differences with respect to neutral stimuli, arguing against unspecific differences between the groups (all *ps*_*bonferroni*_ > 0.05, **Figure. 1B**). ANOVA including group (controls vs. non-dependent vs. dependent) and time (before vs. after cue exposure) and cannabis craving as dependent variable revealed significant main effects for both factors (*F*_(2,79)_ = 44.8, *p* < 0.001, *η*^*2*^ = 0.531; *F*_(1,79)_ = 40.4, *p* < 0.001, *η*^*2*^ = 0.312) and a significant interaction effect (*F*_(2,79)_ = 5.02, *p* < 0.01, *η*^*2*^ = 0.078). Post-hoc comparisons revealed that craving was generally higher in both cannabis groups relative to controls (all *p*_*bonferroni*_ < 0.001). Moreover, within both cannabis groups craving increased after cue-exposure (both *p*_*bonferroni*_ < 0.001, see **Figure. 1C**). Importantly, the cannabis groups did not differ in self-reported trait impulsivity, arousal and craving (all *t* < 1) arguing against confounding effects of these variables on between-group differences with in neural cue reactivity.

**Figure. 1.**
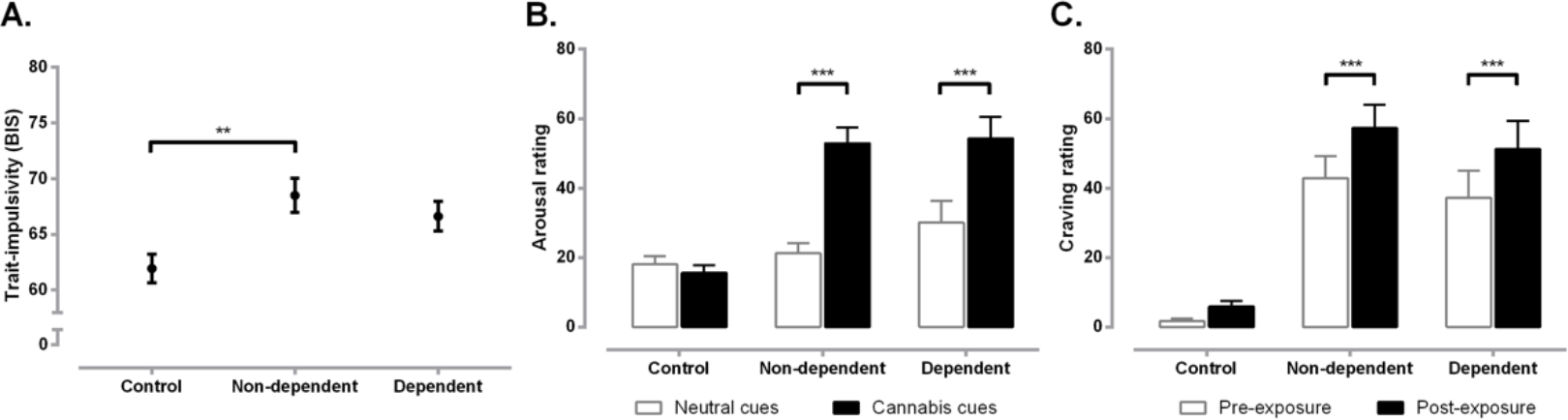
Self-reported trait impulsivity, arousal and cannabis craving in the groups. (A) Both groups of users reported higher trait impulsivity, (B) increased arousal for the cannabis cues, as well as (C) increased cannabis craving after cue-exposure than controls. Mean and standard errors (SEM) are displayed. ** and *** denote relevant significant post-hoc differences at *p*_*bonferroni*_ < 0.01 (A) and *p*_*bonferroni*_ < 0.001 (B and C).

### Neural cue-reactivity - whole brain group differences

Voxel-wise one-way ANOVA (controls vs. dependent vs. non-dependent) of neural cue-reactivity (cannabis > neutral cue) revealed a significant main effect of group predominantly located in the ventral striatum and spreading into the dorsal striatum (see **Figure S1**). Additional effects were observed in a network previously associated with cannabis cue-reactivity (30, 31) encompassing prefrontal, anterior/mid cingulate, superior parietal regions (details **Table S1**). In accordance with the a-priori hypotheses, post-hoc analyses focused on the comparison of cannabis groups with the non-using reference group (see (30, 45, 46) for similar strategy). Relative to controls, non-dependent users demonstrated increased cue-reactivity in a narrow circuit including the ventral striatum (predominantly ventral caudate, spreading into nucleus accumbens), medial prefrontal cortex (MPFC) and right superior parietal cluster (**Figure. 2A**, **Table 3**), whereas dependent users exhibited increased cue-reactivity in a more extensive network, encompassing both, ventral and dorsal striatum as well as limbic, prefrontal, occipital and superior parietal regions (**Figure. 2B** and **Table 3**). These activations largely overlapped with default, dorsal attention, and visual large-scale brain networks (52, 53). Mapping the effects on the atlas-based ventral and dorsal striatal masks further confirmed that both cannabis groups exhibited neural cue-reactivity in the ventral striatum, whereas only dependent users exhibited cue-reactivity in the dorsal striatum (see **Figure. 2C**).

**Table 3.** Brain regions displaying significant cue-reactivity differences between groups

| Cluster region | K (cluster size) | x | y | z | T value |
| --- | --- | --- | --- | --- | --- |
| Non-dependent users > controls |  |  |  |  |  |
| Ventral caudate and NAC<br>extending to MPFC | 152 | -6 | 21 | 9 | 5.33 |
|  |  | -18 | 33 | -9 | 4.80 |
|  |  | -15 | 42 | -9 | 4.50 |
| Superior parietal lobe (precuneus) | 67 | 27 | -54 | 33 | 5.10 |
|  |  | 24 | -42 | 33 |  |
| Dependent users > controls |  |  |  |  |  |
| Limbic lobe extending to temporal,<br>occipital and parietal lobes | 2505 | 27 | -30 | 0 | 6.07 |
|  |  | -18 | -33 | 0 | 5.75 |
|  |  | 6 | 0 | 30 | 5.73 |
| R IFG extending to MFG | 130 | 45 | 9 | 27 | 5.58 |
|  |  | 27 | -9 | 39 | 4.04 |
|  |  | 36 | -6 | 33 | 4.01 |
| L SFG extending to MFG | 112 | -18 | 33 | 42 | 5.37 |
|  |  | -27 | 15 | 42 | 4.00 |
|  |  | -24 | 24 | 45 | 3.55 |
| L IPL extending to PCC/precuneus | 131 | -27 | -60 | 45 | 5.35 |
|  |  | -18 | -51 | 45 | 5.05 |
|  |  | -33 | -51 | 48 | 4.07 |
| Left fusiform | 197 | -42 | -54 | -6 | 5.05 |
|  |  | -30 | -78 | -12 | 4.94 |
|  |  | -45 | -48 | -12 | 4.41 |
| R IFG | 78 | 45 | 30 | 15 | 4.96 |
| MPFC extending to ACC | 200 | -9 | 42 | -3 | 4.67 |
|  |  | -15 | 60 | 9 | 4.36 |
|  |  | -15 | 63 | 18 | 3.93 |
| L IFG extending to MFG | 145 | -45 | 6 | 24 | 4.31 |
|  |  | -39 | 9 | 18 | 4.29 |
|  |  | -33 | -9 | 24 | 3.86 |
| L IFG | 95 | -51 | 39 | 6 | 4.03 |
|  |  | -42 | 33 | 12 | 3.90 |
|  |  | -51 | 30 | 18 | 3.63 |

Note: L =left, R = right, MPFC = medial prefrontal cortex, NAC = nucleus accumbens, ACC = anterior cingulate cortex, IFG = inferior frontal gyrus, MFG = middle frontal gyrus, IPL = inferior parietal lobule, SFG = superior frontal gyrus, PCC = posterior cingulate cortex and dIPu = dorsal lateral putamen. All clusters passed the threshold at cluster level $p_{FWE} < 0.05$ on the whole brain level.

**Figure. 2.**
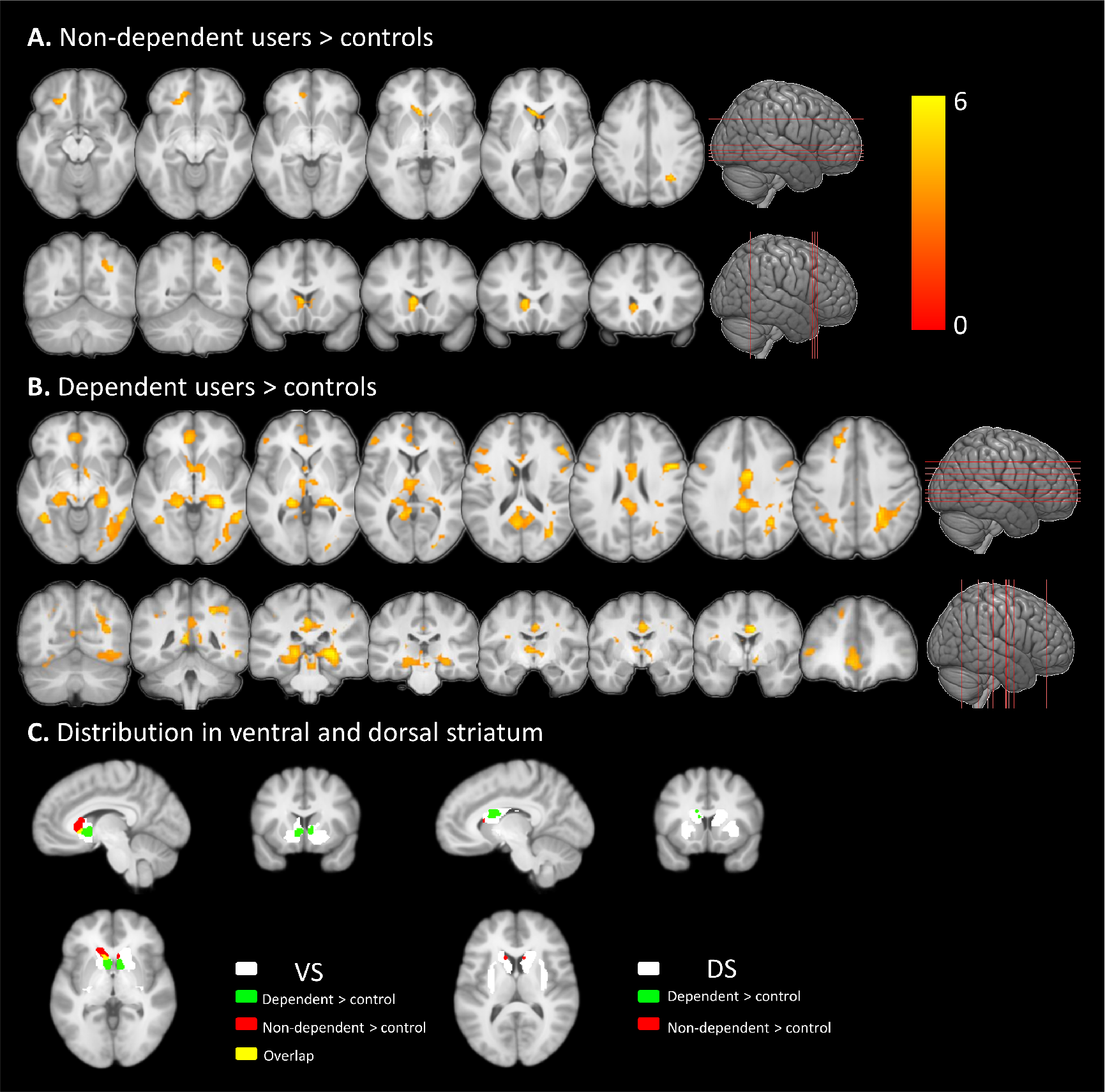
Whole brain cue-reactivity networks in (A) non-dependent, and (B) dependent cannabis users relative to controls. Cue-reactive regions were determined using the contrast [cannabis cues > neutral cues], activation color bars for both groups (displayed in A, B) were scaled to the same range. (C) The activity distribution from (A) and (B) located in ventral and dorsal striatum mask separately. Results displayed at whole-brain cluster level *p*_*FWE*_ < 0.05.

### Striatal subregion-specific contribution to cue-reactivity

To fcharacterize relative contributions of the dorsal vs. ventral striatum (see also (17)), region-specific cue-reactivity estimates were extracted and subjected to mixed ANOVA with group (dependent vs. non-dependent) and subregion (dorsal vs. ventral). Findings revealed a significant main effect of group (*F*_(1, 34)_ = 4.1722, *p* = 0.049; idependent > non-dependent). Post-hoc comparisons revealed that the non-dependent group exhibited significantly elevated ventral compared to dorsal striatal cue-reactivity (*p*_*bonferroni*_ < 0.01, **Figure. 3**), whereas in the dependent group both regions showed comparable high reactivity (both *p*_*bonferroni*_ > 0.05 controlling for abstinence time and age of first use). An additional analysis revealed that both regions exhibited comparably low reactivity to cannabis cues in controls (see plot in **Figure 3**).

**Figure. 3.**
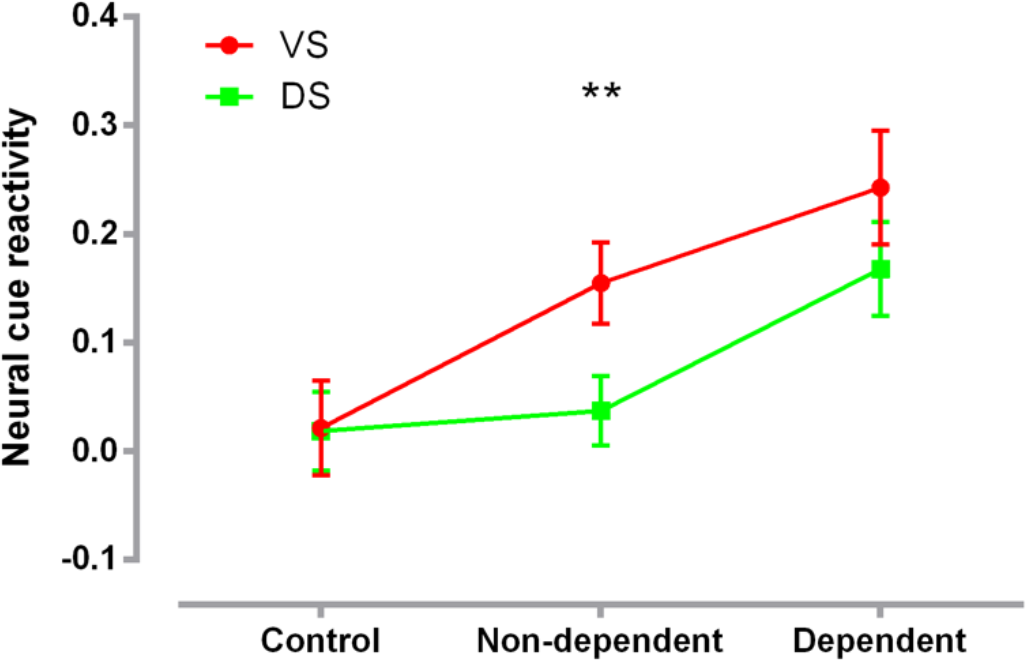
Extracted neural cue-reactivity (cannabis cue > neutral cue) for the ventral (VS) and dorsal (DS) striatum masks. Cue-reactivity in the ventral relative to the dorsal striatum was significantly higher in the non-dependent users, whereas no significant differences were observed in the other groups. Mean and standard errors (SEM) are displayed. ** *p*_*bonferroni*_ < 0.01.

### Striatal network alterations

Non-dependent users exhibited increased dorsal striatal-left medial/superior frontal coupling relative to controls (−30/57/21, *p*_*FWE*_ < 0.05, see **Figure. 4A**), whereas no alterations for ventral striatal networks were observed. In contrast, dependent users exhibited increased dorsal striatal coupling with the left inferior frontal (−36/30/0, *p*_*FWE*_ < 0.05, see **Figure. 4B**) and the ventral anterior cingulate (−9/39/0, *p*_*FWE*_ < 0.05, see **Figure. 4B**) as well as decreased coupling of both, the dorsal and ventral striatum, with a right limbic cluster encompassing hippocampal and amygdala regions (ventral, 36/-15/-21, *p*_*FWE*_ < 0.05, see **Figure. 4C**; dorsal, 36/-18/-24, *p*_*FWE*_ < 0.05, see **Figure. 4D**, details provided in **Table 4**)

**Table 4.** Brain regions of significant functional connectivity differences between groups

| Cluster region | K (cluster size) | x | y | z | T value |
| --- | --- | --- | --- | --- | --- |
| Non-dependent users > controls |  |  |  |  |  |
| DS |  |  |  |  |  |
| L MFG extending to SFG | 61 | -30 | 57 | 21 | 4.14 |
|  |  | -30 | 54 | 30 | 3.95 |
|  |  | -21 | 66 | 15 | 3.72 |
| Dependent users < controls |  |  |  |  |  |
| VS |  |  |  |  |  |
| R Hip extending to Amy | 70 | 36 | -15 | -21 | 4.26 |
|  |  | 36 | 9 | -30 | 3.99 |
|  |  | 30 | 0 | -27 | 3.97 |
| DS |  |  |  |  |  |
| R Hip | 42 | 36 | -18 | -24 | 4.57 |
|  |  | 42 | -27 | -30 | 3.71 |
|  |  | 42 | -12 | -21 | 3.60 |
| Dependent users > controls |  |  |  |  |  |
| DS |  |  |  |  |  |
| L IFG | 122 | -36 | 30 | 0 | 4.80 |
|  |  | -42 | 48 | 6 | 4.40 |
|  |  | -48 | 42 | 3 | 4.13 |
| ACC | 84 | -9 | 39 | 0 | 4.66 |
|  |  | 6 | 60 | 12 | 3.75 |
|  |  | 0 | 54 | 26 | 3.75 |

Note: L =left, R = right, DS = dorsal striatum, VS = ventral striatum, ACC = anterior cingulate cortex, IFG = inferior frontal gyrus, MFG = middle frontal gyrus, SFG = superior frontal gyrus Hip = hippocampus and Amy = amygdala. All clusters passed the threshold at cluster level $p_{FWE} < 0.05$ .

**Figure. 4.**
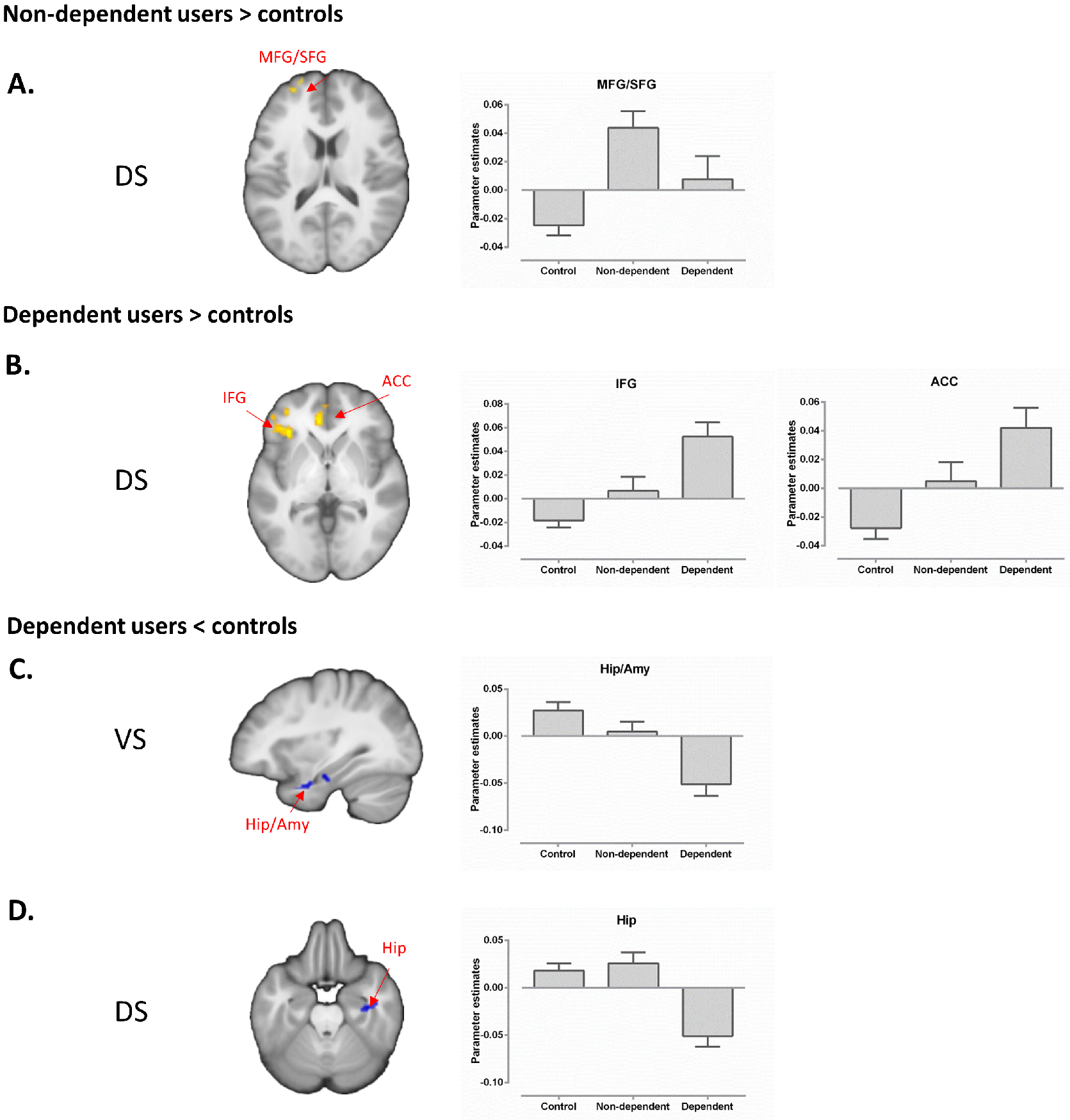
Cue-induced alterations in dorsal and ventral striatal coupling in non-dependent and dependent cannabis users. For visualization, extracted parameter estimates from the target regions are displayed. (A) Non-dependent users exhibited increased dorsal striatal coupling with MFG/SFG, whereas dependent users exhibited (B) increased coupling between dorsal striatum and the IFG and ACC, as well as decreased ventral (C) and dorsal (D) striatal coupling with limbic regions. Results displayed at cluster level *p*_*FWE*_ < 0.05. Abbreviation: DS = dorsal striatum, VS = ventral striatum, Amy = amygdala, Hip = hippocampus, IFG = inferior frontal gyrus, ACC = anterior cingulate cortex, MFG = middle frontal gyrus and SFG = superior frontal gyrus.

**Figure. 5.**
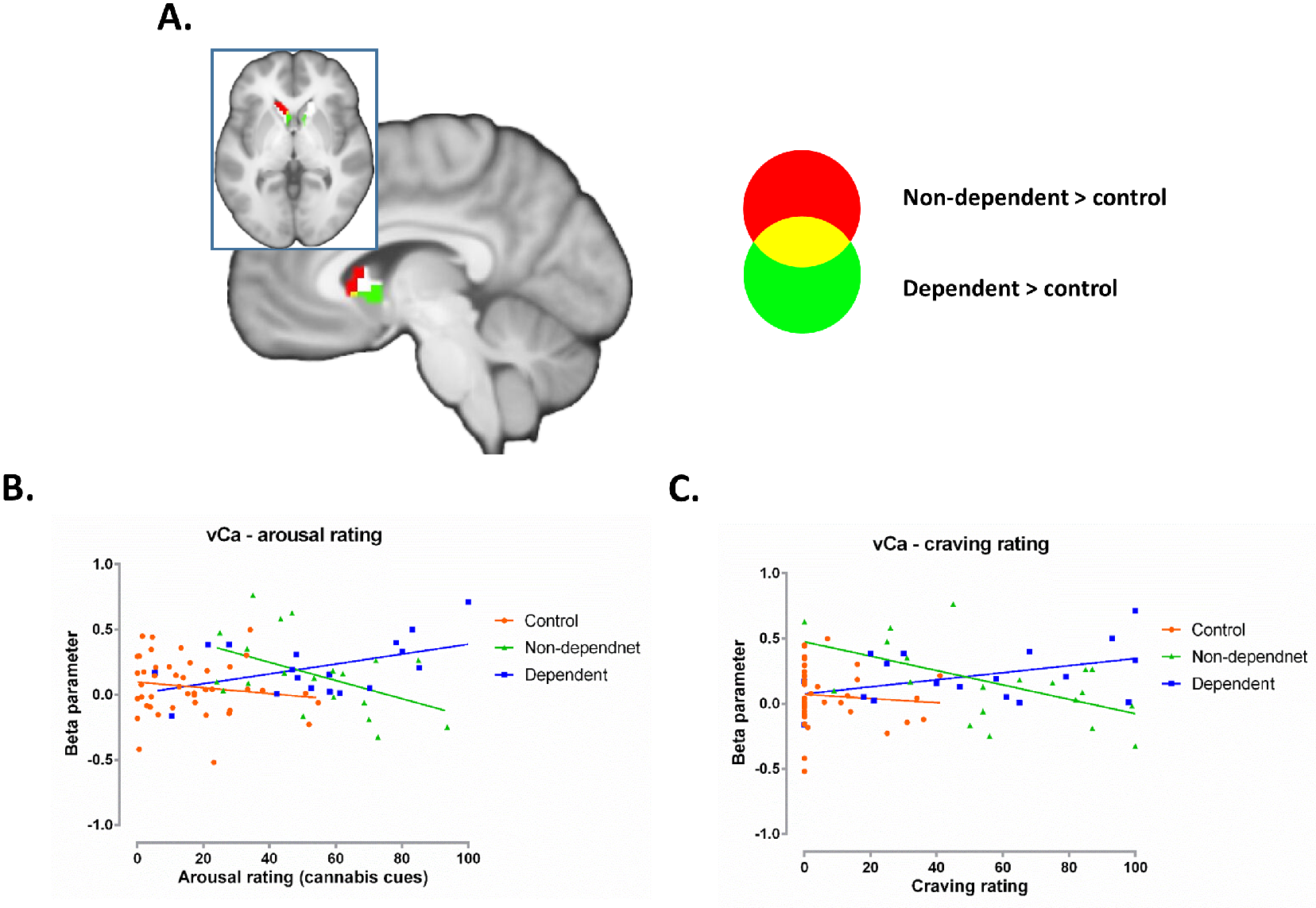
(A) displays overlapping voxels located in the ventral caudate (vCa, red: non-dependent > control, green: dependent > control, yellow: overlap, white: vCa mask). Ventral caudate cue-reactivity was positively associated with both cue-induced arousal (B) and craving (C) in dependent users. In non-dependent users, ventral caudate cue-reactivity was negatively correlates with arousal and craving.

### Brain behavior associations

Given that common cue-reactivity alterations were predominantly observed for the ventral caudate, parameter estimates were extracted from this region and entered into correlation analyses. Ventral caudate cue-reactivity was positively associated with cue-induced arousal in dependent users (*r* = 0.469, *p* = 0.0496), but negatively correlated in non-dependent users (*r* = −0.478, *p* = 0.033, significant between group correlation differences; *z* = 2.91, *p* = 0.0037; **Figure. 5B**). A (marginally) significant positive association was found between neural cue-reactivity and post cue-exposure craving ratings in dependent users (*r* = 0.433, *p* = 0.073), but a significant negative correlation in non-dependent users (*r* = −0.559, *p* = 0.010, *z* = 3.091, *p* = 0.002, see **Figure. 5C**). No significant correlations were observed in controls (arousal *p* = 0.325; craving *p* = 0.593)

### Additional control analysis

No between-group differences with respect to brain structure were observed in whole brain or striatum-focused analyses arguing against confounding effects of brain structural alterations on dependence-group specific cue-reactivity (details provided in **Supplementary Materials**). Age of onset or time since last use were not significantly associated with cue-reactivity indices - including VS and DS-specific reactivity - arguing against strong confounding effects (all p > .17, in separate and pooled cannabis groups).

## Discussion

The present study aimed at determining neural cue-reactivity that specifically characterizes cannabis addiction while accounting for adaptations associated with cannabis exposure. To this end, cannabis users were stratified according to their dependence status (dependent and non-dependent) and compared to carefully matched non-using controls. As expected, cue-exposure increased arousal and craving and elicited exaggerated neural reactivity in regions previously associated with drug cue-reactivity irrespective of the dependence status. In line with our hypotheses both groups of cannabis users exhibited exaggerated ventral striatal reactivity in response to cannabis-cues and elevated trait impulsivity, whereas dorsal striatal cue-reactivity was specifically observed in dependent users. On the network level, both groups of cannabis users demonstrated increased dorsal striatal-prefrontal coupling, whereas dependent users additionally exhibited decreased coupling of both striatal sub-regions with limbic regions encompassing the right hippocampus and amygdala. Exploratory analyses further revealed that the level of cue-induced arousal and craving correlated negatively with ventral caudate reactivity in non-dependent cannabis users and that this association was reversed in dependent users.

In line with our region-specific hypothesis cannabis cue-exposure produced exaggerated ventral striatal activity in both groups of cannabis users relative to non-using controls. The ventral striatum is strongly engaged in signaling reward value and anticipation and thus contributes to the incentive salience of drug cues as well as associated decision making, including impulsive behavior (40, 41, 54,55). Ventral striatal cue-reactivity has been consistently observed in meta-analytic studies covering data from frequent users of different classes of drugs (26–29) and has been considered to reflect the exaggerated salience of drug-associated stimuli. In contrast, ventral striatal reactivity towards non-drug rewards has been frequently found blunted in drug using populations, including cannabis users (56) and may represent a predisposing vulnerability for escalating substance use (57) as well as a consequence of chronic cannabis exposure (58). In support of the present findings, recent studies reported marked reward- and salience-related electrophysiological responses to drug cues across infrequent and heavy cannabis users (59) suggesting that an incentive sensitization of the ventral striatal reward system may promote but not fully explain the transition to addictive cannabis use (30, 60, 61). High levels of impulsivity have been frequently observed across heavy drug users as well as their biological relatives (62) and individual variations in this trait have been linked to ventral striatal dopamine function (40, 41). Translational models suggest that the increased vulnerability to escalate drug intake and develop compulsive use in rodents with high impulsivity is mediated by the ventral striatum (63, 64). Increased impulsivity in both cannabis using groups may therefore suggest that elevated levels of trait impulsivity increase the propensity to escalate substance use which may in turn be further exacerbated by chronic cannabis exposure (for similar findings on trait impulsivity in stimulant addiction see also (39, 62)).

In contrast, exaggerated dorsal striatal cue-reactivity was selectively observed in dependent users, suggesting an important contribution of the dorsal striatal subregion to cannabis addiction. Whereas the ventral striatum is critically involved in salience signaling and initial learning of goal-directed behavior (65), the dorsal striatum critically mediates the transition to habitual, stimulus-controlled behavior (66). In line with the functional differentiation of the striatum, animal models of addiction suggest that the dorsal striatum controls the progression from goal-directed to cue-controlled drug seeking and taking (5, 51). Whereas the shift between ventral to dorsal striatal control over behavior has been extensively demonstrated in laboratory animals only two studies explored whether these findings translate to the human condition. Combining cue-exposure with neuroimaging these studies demonstrated that drug-cues elicit neural reactivity in the dorsal striatum of heavy but not light alcohol drinkers (35), and craving-associated dopamine release in the dorsal but not ventral striatum in cocaine-dependent individuals (67). Previous studies probing neural cue-reactivity in cannabis users provided some indirect evidence that responses in ventral striatal reward pathways may reflect exposure-related adaptations, whereas adaptations in the dorsal striatum may mediate addictive processes (30, 31), including habitual drug seeking (34). Together the present findings resonate with these previous reports and suggest that adaptations in the dorsal striatum mediate the transition to dependent cannabis use in humans.

On the whole brain level, dependent cannabis users exhibited neural cue-reactivity in a widespread network encompassing frontal, occipital, limbic, temporal and superior parietal regions whereas non-dependent users exhibited more focal increases in medial prefrontal and superior parietal regions. Abnormal cue-reactivity in these regions has been reported in previous studies in heavy cannabis users (30, 61, 68), with the present findings suggesting that addiction-related neuroadaptations are not specifically limited to the dorsal striatum. From a network-perspective the widespread hyperactive network observed in dependent users encompasses core regions of the default mode network, including posterior cingulate cortex/precuneus, medial prefrontal, hippocampal and parietal regions, which plays an important role in the evaluation of self-related and highly salient information (69). Greater activation in dependent cannabis users may thus reflect exaggerated salience attributed to drug cues, which in turn may promote drug seeking.

At the network level, cannabis users exhibited increased cue-induced dorsal striatal communication with prefrontal regions regardless of dependence status. Aberrant intrinsic and task-based striatal communication with frontal regions engaged in reward processing and regulatory control has been repeatedly reported in cannabis users (17, 70–72). The fronto-striatal circuits are engaged in several addiction-relevant functional domains including incentive salience processing and flexible behavioral control (1, 3, 36) and alterations in this circuitry may reflect exaggerated salience of drug-associated cues (71) or deficient regulatory control over behavior (17). Cannabis dependent participants additionally demonstrated decreased connectivity of both striatal subregions with limbic regions encompassing the hippocampus and amygdala. Both regions are at the core of emotional memory formation with the amygdala mediating the impact of emotional experience on contextual memory formation in the hippocampus. During the transition to addiction both regions are thought to interact with the striatum to establish the impact of drug-associated cues on habitual behavior (2). Drug exposure is considered to promote habitual drug seeking behavior while suppressing processing of other information (28, 51) resulting in a biased evaluation (73) and an increased motivational drive to use the drug. Together the present network level findings may thus suggest that both groups of cannabis users exhibit increased salience signaling and deficient frontal regulatory control while subcortical emotional memory circuits involved in habitual behavior are specifically dysfunctional in dependent users.

Finally, an exploratory analysis revealed divergent associations between ventral caudate cue-reactivity and the degree of cue-induced craving and arousal in dependent and non-dependent cannabis users. However, these findings need to be interpreted with caution due to the exploratory nature of this correlational analysis. Previous research using cue-exposure paradigms reported that higher levels of arousal and craving are linked with stronger ventral striatal cue reactivity (28, 74, 75). Whereas dependent cannabis users in the present study resembled this association, non-dependent users demonstrated the opposite pattern, which may reflect a protective mechanism possibly related to intact regulatory control over cue-induced craving.

## Limitations and conclusion

Although the present study design allowed to control for important confounders including the co-use of other drugs and alterations associated with chronic exposure to cannabis, findings need to be considered in the context of the following limitations: (1) although evidence from animal models indicates that the ventral and dorsal striatum are differentially impacted during the progression to addiction, cross-sectional studies in humans are not sufficient to allow causal inferences in humans which can only be established by prospective longitudinal designs, (2) the present study focused on male cannabis users and an increasing number of studies reported differential effects of cannabis in male and female users (76). Future research thus needs to determine whether the observed findings generalize to cannabis-dependent women.

Together, the present study demonstrated common and distinguishable neural reactivity towards drug-associated cues in dependent and non-dependent users. Both groups showed increased ventral striatal reactivity and striato-frontal connectivity possibly reflecting exaggerated salience of drug cues, whereas increased dorsal striatal and suppressed striato-limbic connectivity was only evident in dependent users possibly reflecting neuroadaptations in circuits underlying habitual responses and compulsive drug seeking.

## Acknowledgements

We thank the authors of previous cannabis cue-reactivity studies (30, 68) for providing the cannabis-associated stimuli.

The authors declare no competing interests.

