## Supplemental for "Cue-reactivity in the ventral striatum characterizes heavy cannabis use, whereas reactivity in the dorsal striatum mediates dependent use"

### Supplementary information

**Zhou et al.,**

Contact:

### Supplementary Materials and Methods

#### Participants

Inclusion criteria for all participants were right-handedness and age 18-35 years. Exclusion criteria for all participants included 1) history of DSM-IV disorders, such as psychotic or bipolar symptoms (validated by the MINI), 2) increased levels of depression (Beck Depression Inventory, BDI-II, score > 20, (1)), 3) medical disorder, 4) current or regular medication, 5) usage of other illicit substances >60 lifetime occasions or during the 28 days prior to the experiment, 6) >20 cigarettes per day, 7) positive urine screen for cocaine (300 ng/ml), methamphetamine (500 ng/ml), amphetamine (500 ng/m), methadone (300 ng/ml) or opiate (300 ng/ml) (Drug-Screen-Multi 7TF, nal von minden GmbH, Moers, Germany), 8) breath alcohol level >0.00‰ (assessed by breath sample using TM-7500, Trendmedic, Penzberg, Germany). Controls were included if their cumulative lifetime use of cannabis was < 15 g. Additional exclusion criteria for controls were use of any other illicit substance >10 lifetime occasions. To control for confounding acute effects of cannabis, all users had to remain abstinent 24 h prior to the experiment. To promote adherence to abstinence, participants were informed that recent use would be detectable by the urinary drug tests on the day of the experiment (see similar procedures in (2, 3), for the use of a bogus pipeline to maintain abstinence in cannabis users see also (4)). Trait impulsivity was assessed by means of the Barratt Impulsiveness Scale (BIS (5)). To further control for potential confounding effects of between group differences in attention and state anxiety, the State Anxiety Inventory (STAI (6)) and the d2 test of attention (7) were administered on the day of the experiment.

**Experimental protocols and cue-exposure procedure**

During the cue-reactivity paradigm, participants were shown nine blocks depicting either cannabis-cue or neutral stimuli (5 stimuli per block, each stimulus presented 3 s, inter-trial interval 0.5 – 1.5 s, inter-block interval 14.5 – 15.5 s). Cannabis-cue stimuli were validated in previous studies on cue-reactivity in cannabis users (8, 9), neutral stimuli were selected from the IAPS and NAPS databases (10, 11). To ensure attentive processing, each block included a randomly selected picture with a red frame that required a button press from the subjects. After each block, a visual analog scale (VAS, ranging from 0-100%; randomized starting point between 25% ~ 75%) was presented for 5s to assess cue-induced increases in physiological arousal (12). We opted for a rating of arousal instead of craving because arousal may allow a more suitable comparison with controls given that they never have experienced actual cannabis craving. To assess cue-induced craving, cannabis craving was assessed directly before and after the paradigm in the MRI scanner (VAS, 0-100% cannabis craving). All responses had to be confirmed by a second button press which was visually indicated to the subjects by a change in the color of the scale. To control order effects, the order of blocks and the order of stimuli within each block were randomized. Paradigms were presented using Presentation 14 (Neurobehavioral Systems, Albany, CA) and liquid crystal display video goggles (Nordic NeuroLab, Bergen, Norway).

### fMRI acquisition, processing and first level model

Data was acquired on a Siemens Trio 3T MRI system (Siemens, Erlangen, Germany). Functional data was acquired using a T2* echo-planar imaging (EPI) BOLD sequence (repetition time (TR) = 2500 ms, echo time (TE) = 30 ms, 37 slices, voxel size = 2 × 2 × 3 mm, flip angle = 90°, field of view = 192 mm). To exclude subjects with apparent brain pathologies and facilitate normalization of the functional data, a high-resolution T1-weighted structural image was additionally acquired (TR = 1660 ms, TE = 2.54 ms in 208 slices, field of view = 256 mm, voxel size = 0.8 × 0.8 × 0.8 mm).

MRI data was processed using SPM12 (Welcome Department of Cognitive Neurology, London, UK, http://www.fil.ion.ucl.ac. uk/spm/software/spm12) (13) implemented in MATLAB Version 8.3 (Math Works Inc., Natick, MA). The time series were initially realigned to correct for head movement based on a six-parameter rigid body algorithm. T1 and functional images were co-registered and the functional timeseries were subsequently normalized to MNI space with a 3 × 3 × 3 mm voxel resolution applying normalization parameters determed by segmenting the T1 images. The normalized data were subsequently spatially smoothed with a Gaussian kernel at full width at half maximum (FWHM) of 6 × 6 × 6 mm. Participants were excluded from further analysis, if they displayed excessive head motion (> 3 mm or > 3°). Session-specific regressors for the experimental conditions (neutral, drug cues) and the rating periods following each block were modelled in the first level design matrix and convolved with the canonical hemodynamic response function. The six realignment parameters were included as regressors to improve motion control. In line with the main aim of the study, the subsequent group-level analysis was carried out using the first-level contrast determining neural cue-reactivity (cannabis cue > neutral cue).

**Task-modulated functional connectivity**

Based on our a priori regional hypotheses the VS and DS masks were employed as seed regions for the gPPI analysis. White matter and cerebrospinal fluid signals were controlled for by linear regression and linear detrending. Extracted BOLD time series from the subregion-specific masks were treated as main physiological factors in the gPPI design matrix. Experimental conditions and contrasts of interest were defined in line with the BOLD level analysis.

**VBM analysis and results**

To further control for potential confounding effects of brain structural differences between the cannabis using groups, an additional voxel-based morphometry (VBM) analysis was carried out using the T1-weight MRI data. Standard protocols for VBM analyses as implemented in the CAT12 toolbox (http://dbm.neuro.uni-jena.de/cat/) were employed. Briefly, the spatial normalization was used to register the MRI images to the standard Montreal Neurological Institute (MNI) space with affine registration. The tissue segmentation was used to classify data into gray matter (GM), white matter (WM) and cerebrospinal fluid (CSF) with modulated normalization. The total intracranial volume (TIV) was included as covariate to control for individual brain size. Before statistical inference, the segmented images were convolved with an isotropic Gaussian Kernel (8 mm) to increase the signal-to-noise ratio.

After the preprocessing steps concordant GLM ANOVA models as for the functional data were employed on the second level. No differences were observed in this exploratory analysis (whole brain or striatum search volume).

**Table S1 Main effect of group in cue reactivity.**

| Cluster region | K (cluster size) | x | y | z | Z value |
|---|---|---|---|---|---|
| R Hip | 149 | 27 | -30 | 0 | 5.08 |
| | | 27 | -15 | -9 | 3.14 |
| Parietal lobel | 253 | 27 | -54 | 33 | 5.02 |
| | | 24 | -45 | 42 | 4.52 |
| | | 30 | -54 | 45 | 4.42 |
| R MCC | 216 | 6 | 0 | 30 | 5.00 |
| | | -3 | 9 | 24 | 4.64 |
| | | -6 | 21 | 9 | 4.55 |
| L Hip | 97 | -18 | -33 | 0 | 4.86 |
| | | -12 | -24 | -9 | 3.27 |
| L Precuneus | 246 | -15 | -48 | 9 | 4.79 |
| | | -3 | -57 | 15 | 4.37 |
| | | -3 | -33 | 30 | 4.31 |
| R ITG | 82 | 48 | -54 | -6 | 4.68 |
| R IFG | 57 | 45 | 9 | 27 | 4.61 |
| L ACC | 101 | -21 | 33 | -12 | 4.58 |
| | | -9 | 42 | -3 | 4.13 |
| L SPG | 50 | -27 | -60 | 45 | 4.41 |
| | | -18 | -51 | 45 | 4.33 |
| | | -33 | -51 | 48 | 3.37 |
| Temporal lobe | 51 | -42 | -54 | -6 | 4.26 |
| | | -45 | -48 | -12 | 3.79 |
| L fusiform | 54 | -30 | -78 | -12 | 4.24 |
| | | -24 | -78 | -6 | 3.59 |
| | | -18 | -84 | -12 | 3.31 |
| R fusiform | 111 | 33 | -51 | -18 | 4.19 |
| | | 36 | -66 | -6 | 3.81 |

Note: L =left, R = right, ACC = anterior cingulate cortex, MCC = middle cingulate cortex, IFG = inferior frontal gyrus, Hip = hippocampus, ITG = inferior temporal gyrus, and SPG = superior parietal gyrus. All clusters passed the threshold at uncorrected $p$ = 0.001 and cluster level

$p_{FWE} < 0.05$ for whole brain results.

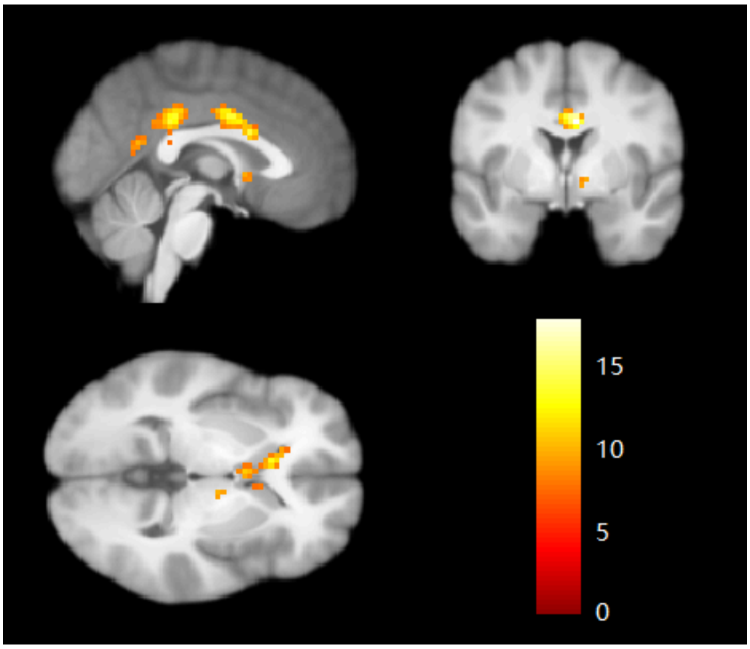


**Figure. S1** Main effect of group displayed at voxel-level uncorrected ($p < 0.001$) and cluster k = 200 (for display purpose)

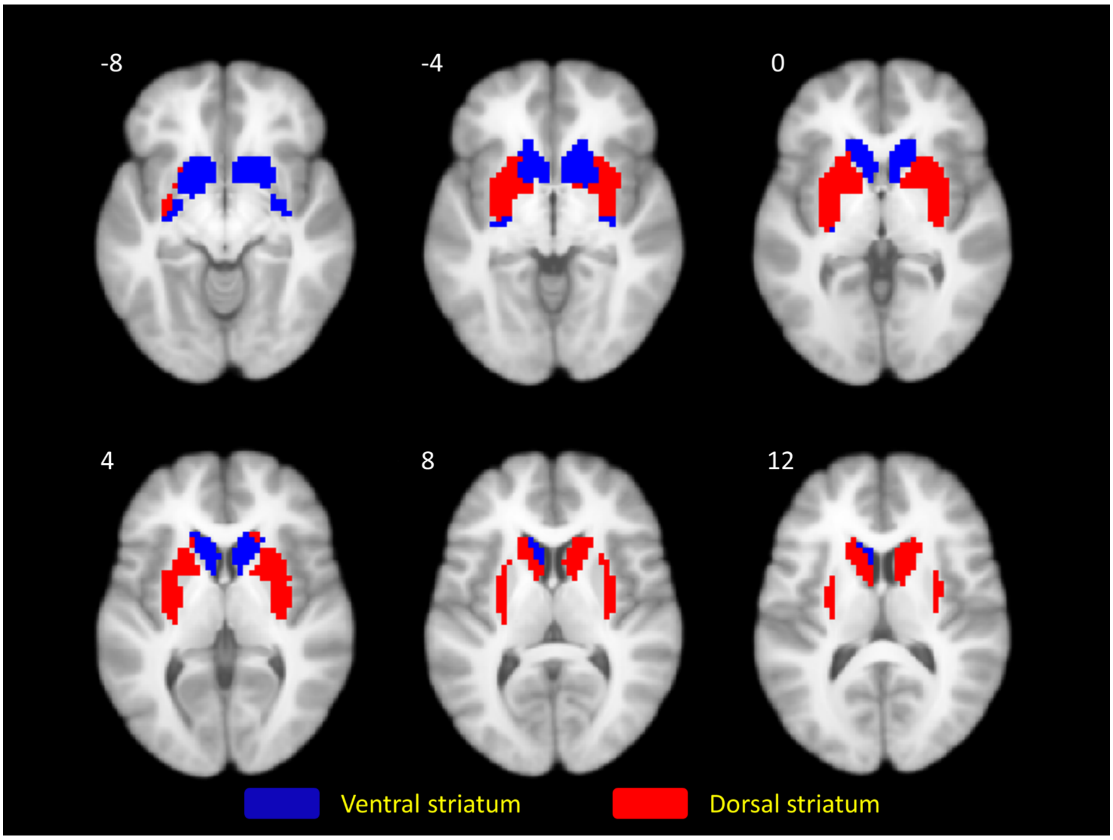


**Figure. S2** Anatomical masks, i.e. the ventral striatum mask (blue) and dorsal striatum mask (red), resampled at 3mm. The number indicates slice locating in the z axis.
